# A wild genome of the underutilized legume lablab (*Lablab purpureus*) reveals the genetic basis of domestication

**DOI:** 10.1101/2025.11.13.688277

**Authors:** Tsz-Yan Cheung, Delbert Almerick Tan Boncan, Yuxuan Yuan, Yunjia Zhang, Prakit Somta, Mark A. Chapman, Ting-Fung Chan

**Author notes:** Corresponding author (Ting-Fung Chan).

## Abstract

**Background:** With global food security threatened by climate change and the growing global population, underutilized crops offer sustainable solutions for diversifying current agricultural systems. Lablab (*Lablab purpureus*) is a highly versatile, climate-resilient underutilized legume native to Africa and widely, but locally, cultivated in Asia with untapped potential for sustainable agriculture. However, the lack of wild genome resource prevents a comprehensive genome utilization research for this underutilized crop. This paper presents the first chromosome-scale genome of wild lablab with its complete gene annotation and provides comprehensive population genomic analyses of its evolution and domestication by resequencing a panel of wild and domesticated accessions.

**Results:** The genome was assembled into a total size of 478.2 Mb with a scaffold N50 of 39.7 Mb, and 96.4% (460.8 Mb) of the scaffolds anchored to 11 pseudochromosomes, the haploid chromosome number of lablab. We demonstrated that domestication was likely in Africa with several subsequent instances of transportation into Asia. Our analyses identified selective sweeps resulting from domestication as well as pseudogenes associated with yield-related traits and stress tolerance, representing candidate genes for further validation and targets for future breeding programs.

**Conclusions:** Our genomic resource for wild lablab delineated its domestication history and identified key genes and pseudogenes under selection, bridging the gap between its untapped wild diversity and breeding for improved stress resilience and yield in underutilized legumes. This offers genomic and genetic insights for crop improvement of lablab.

## Background

The rising global population, accompanied by an increasing food demand, poses a major threat to food security. Meanwhile, breeding staple crops remains challenging amid recent climate events, which have led to a significant loss of yield of several major crops due to elevated global temperature [1, 2]. To counter the adverse effects of climate change on the yield of existing crops, cultivating underutilized crops represents a promising approach to enhance food security [3]. Underutilized crops, also known as neglected or orphan crops, are indigenous plant species that are well adapted to specific ecological niches. This adaptation is conferred by agronomic traits, such as stress tolerance or resistance, whose selective advantages allow the crops to survive and thrive. Therefore, these underutilized crops offer the potential to address the threats posed by climate change to food security by improving and augmenting crop production [4, 5].

*Lablab purpureus* (L.) Sweet (widely known as lablab or hyacinth bean), the sole species in the *Lablab* genus, is an underutilized legume from Africa with over 3,000 reported accessions in the tropical and subtropical regions [6, 7]. Based on the number of seeds per pod, lablab can be broadly categorized as two- and four-seeded. This dichotomy reflects two distinct gene pools, with domesticated populations derived from within both [8–10]. Within the four-seeded domesticates is a subset of accessions with up to eight seeds per pod named ssp. *bengalensis.* Despite the wild species being native to Africa, domesticated lablab is most widely grown in South-East Asia.

Lablab is an agro-morphologically and physiologically diverse legume with greater drought tolerance than other legumes, such as *Vigna unguiculata* (L.) Walp. (cowpea) and *Phaseolus vulgaris* L. (common bean) [6, 11, 12]. It is rich in nutrients, including proteins, vitamins, minerals, essential omega-3, and omega-6 fatty acids [13], that could be leveraged to alleviate malnutrition and enhance nutritional security. Moreover, it is a versatile species whose parts can be utilized for several purposes. Its pods, seeds, and leaves are consumed as vegetables, dried white seeds of specific cultivars are used as a Chinese herbal medicine [14], and seeds with a high level of lauric acid could be used as nutraceuticals [15]. Although there is an apparent incentive to cultivate lablab, its potential as an economic crop has yet to be realized. An in-depth understanding of the genetic diversity of lablab is essential for us to utilize it to its full potential.

Recently, genome-based approaches have identified lablab with desired agronomic traits and investigated the potential for marker-assisted breeding [16–18]. Two studies have caried out association analyses, using genotyping-by-sequencing [8] and whole genome sequencing [16] for lablab, identifying several marker-trait associations for agronomic traits, however a ‘bottom-up’ selection analysis [19] identifying genes associated with domestication is lacking. Further, the whole genome population genomic analysis of Teshome *et al*. [16] suggested that cultivated accessions did not group by geography, and in at least once case wild two-seeded, domesticated two-seeded and wild four-seeded accessions formed a group, which conflicts with the established taxonomy. The geography of lablab domestication and the relationship between geographic and genetic variation is therefore still unclear.

Incorporating lablab into the current food system outside where it is currently grown and eaten, requires a comprehensive assessment of its genetic diversity to facilitate the development of varieties with desired traits, such as high yield and stress resilience. Considering that successful genetic improvement of crops depends on available genome resources, it is imperative to investigate the genome and population genomics of lablab. Moreover, a high-quality reference genome with accurate annotations is crucial for genome-assisted breeding. This need has been met with rapid advancements and cost reduction in sequencing technologies, fuelling a momentum in crop genomics. Advances in genomics have facilitated the study of previously untapped species in the wild whose genetic repertories could be leveraged for selective breeding, and the identification of relevant stress-associated genes in existing cultivars could be utilized to develop cultivars with improved traits [20–22]. To date, several lablab genomes from four-seeded cultivars have been reported, with qualities ranging from draft genomes to chromosome-level assemblies [8, 23–25]. However, a wild reference genome is still lacking, preventing comprehensive genome utilization research for this underutilized crop.

In this study, we generated the first high-quality chromosome-level genome assemblies of wild lablab (*L. purpureus* ssp. *uncinatus*) from the four-seeded gene pool, i.e. the ancestor of the vast majority of domesticated lines. By resequencing a panel of wild and domesticated accessions, we studied the population structure of this underutilized crop, identified a core set of SNPs to differentiate all domesticated samples, and revealed genes positively selected together with pseudogenes during domestication. These findings uncover the molecular and genetic basis of important agronomic traits, thereby providing not only genomic resources but also important insights into the crop’s genetic diversity and its potential for breeding and improvement.

## Results

### Genome assembly

Multiple wild and domesticated lablab accessions were selected for this study (Fig 1a). To assemble a high-quality genome of the four-seeded wild lablab accession (ILRI accession 24779 from Zimbabwe), we generated 1.99 M PacBio HiFi reads (36.27 Gb) with an N50 of 18.1 kb, representing ∼85.7× coverage of the genome based on the previously estimated genome size of 423 Mb [23]. The initial genome was assembled into 189 contigs, with a total size of 500.9 Mb and a contig N50 of 29.5 Mb. To construct a chromosome-level assembly, optical genome mapping (OGM) and Hi-C (Omni-C) were used for scaffolding. Optical mapping molecules of 150 kb or longer with at least nine DLE-1 labels were used. A total of 1,188.8 Gb optical mapping data were generated, yielding a molecule N50 of 235.3 kb. The mapping rate to draft contig-level genome assembly was 50.9%, achieving a final coverage of 1,253.3×. The majority of contigs were scaffolded (i.e., hybrid scaffolding) by the optical genome maps, resulting in an assembly comprising 39 scaffolds, with a total length of 477.8 Mb and an N50 of 21.0 Mb. Although OGM improved the number of contigs, it resulted in a shorter N50 with 729 new gaps. Following hybrid scaffolding, 28.7 Gb of Omni-C data were added in the second round of scaffolding, polishing the genome by correcting the scaffolds and improving the scaffold N50. At this stage, the scaffold-level assembly comprised 859 scaffolds, with a total length of 478.1 Mb and an N50 of 31.9 Mb. The genome was further polished using a reference-based strategy [8], successfully anchoring the scaffolds to 11 pseudomolecules. After gap filling, the final four-seeded wild lablab genome comprised 231 scaffolds, with a total length of 478.2 Mb and an N50 of 39.7 Mb. In total, 96.4% (460.8 Mb) of the scaffolds in the wild lablab genome was anchored to 11 pseudochromosomes (Fig 1b), corresponding to the haploid chromosome number of lablab.

**Fig. 1.**
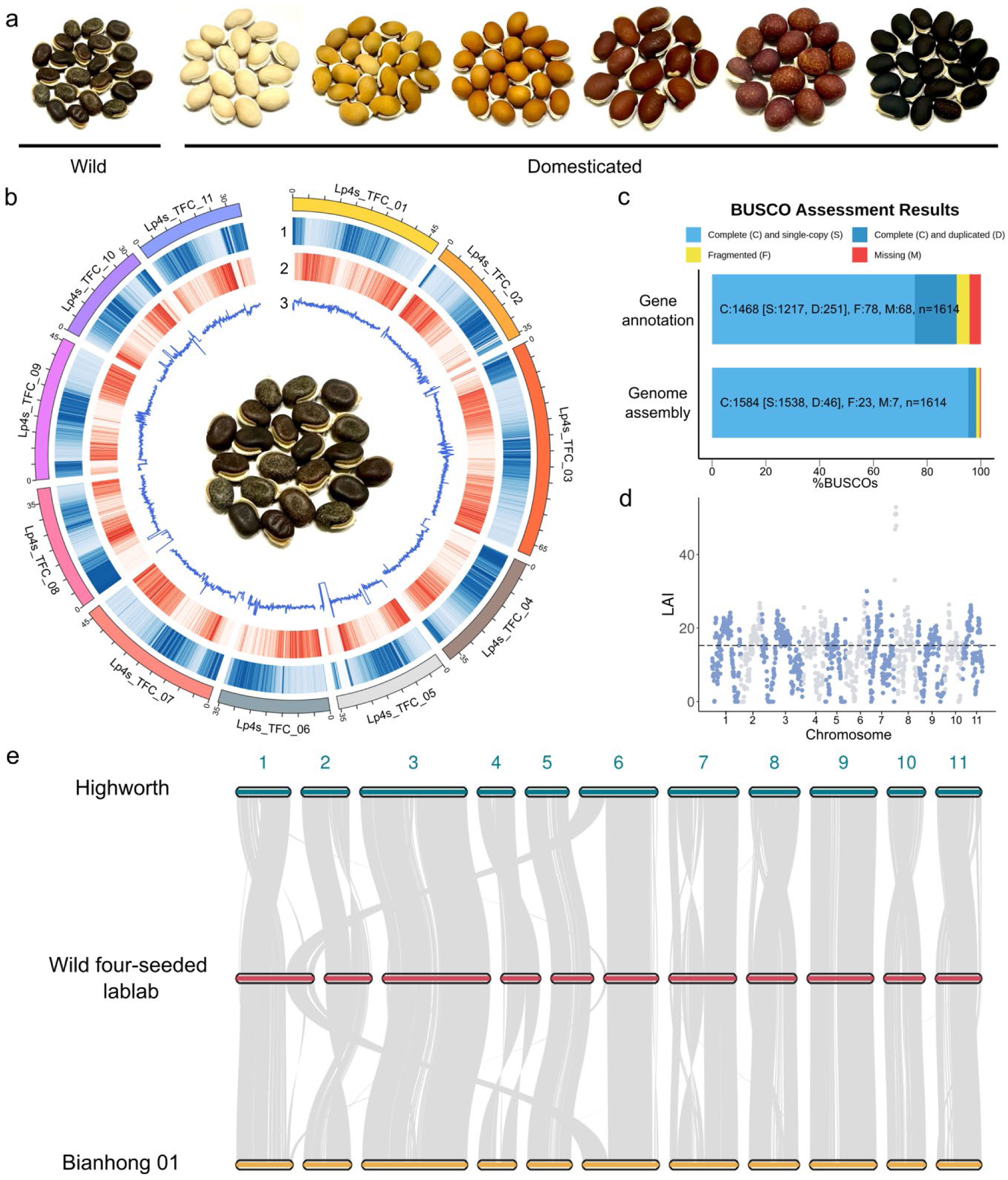
Genome assembly of four-seeded wild lablab. (a) Seeds of some selected *Lablab purpureus* accessions in this study (photos by T.-Y. C.). (b) Distribution of the genomic features of the four-seeded wild genome. The outer layer illustrates the 11 pseudochromosomes with the lengths in megabases (Mb). The outer to inner tracks represent 1) repeat density, 2) gene density, and 3) GC content, all calculated within a 100-kb window. (c) BUSCO assessment of the genome assembly and gene annotation using the embryophyta lineage. (d) LAI distribution of the 11 pseudochromosomes. The dashed horizontal line represents the average LAI value of the genome. (e) Syntenic comparison of the 11 pseudochromosomes between four-seeded wild lablab, cv. Highworth and Bianhong 01.

A Benchmarking Universal Single-Copy Orthologs (BUSCO) assessment illustrated a high level of genome completeness, in which 98.2% of the embryophyta BUSCOs (95.3% as single-copy) were complete in the genome assembly (Fig. 1c; Table 1). Additionally, a whole-genome long terminal repeat (LTR) assembly index (LAI) of 15.28 was obtained for the wild lablab genome, demonstrating a reference-grade genome assembly [26] (Fig. 1d). To further evaluate the completeness of the genome assembly, the 11 pseudochromosomes were compared against the published genomes of lablab cultivars Highworth [8] and Bianhong 01 [25], which have an assembly size of 417.9 Mb and 463.3 Mb respectively. Despite the substantial difference in genome size between the three lablab genomes, the syntenic comparison displayed large blocks of homology between their pseudochromosomes with several chromosomal rearrangements. Notably, there was one inter-chromosomal translocation between chromosome 1 of four-seeded wild lablab and chromosome 6 of two cultivars, proposing the difference in genomic structure between wild and domesticated accessions in lablab (Fig. 1e). Most of the chromosomes displayed complete synteny, while no missing large-scale synteny blocks were found in the three genomes, suggesting that the core genomic structures were preserved at a high degree of completeness.

**Table 1.** Summary of reference genome assembly and annotation.

| Category | Metric | *Lablab purpureus* ssp. *uncinatus* |
| --- | --- | --- |
| Assembly | Accession | 24779 |
|  | Heterozygosity | 0.28% |
|  | Assembly size (bp) | 478,184,590 |
|  | No. of contigs | 189 |
|  | Contig N50 (bp) | 29,508,882 |
|  | No. of scaffolds | 230 |
|  | Scaffold N50 (bp) | 39,682,417 |
|  | L90 | 11 |
|  | Longest scaffold (bp) | 68,650,878 |
|  | GC content | 30.43% |
|  | No. of gaps | 730 |
|  | No. of N in bp (% of genome) | 3,983,856 (0.83%) |
|  | Complete BUSCOs | 98.2% (S: 95.3%, D: 2.9%) |
|  | Average LAI | 15.28 |
| **Gene**<br>**annotation** | No. of annotated genes | 29,004 |
|  | No. of total transcripts | 34,454 |
|  | No. of coding transcripts | 32,756 |
|  | No. of tRNAs | 545 |
|  | No. of lncRNAs | 1,698 |
|  | Complete BUSCOs | 91.0% (S: 75.4%, D: 15.6%) |
| **Repeat**<br>**elements** | LTR in bp (% of genome) | 97,252,800 (20.34%) |
|  | LINE in bp (% of genome) | 1,901,384 (0.40%) |
|  | DNA transposons in bp (% of genome) | 10,693,548 (2.24%) |
|  | Rolling circles in bp (% of genome) | 1,584,263 (0.33%) |
|  | Others in bp (% of genome) | 19,767,684 (4.13%) |
|  | Unclassified repeats in bp (% of genome) | 137,115,063 (28.67%) |
|  | Total repeats in bp (% of genome) | 268,314,742 (56.11%) |

### Genome annotation

A total of 439,878 repetitive sequences were identified in the assembled genome, spanning approximately 268 Mb and accounting for 56.11% of the genome (Table 1). These values are greater than those in the assemblies of cv. Highworth [8] (268,915 repeat sequences covering 43.4% of the genome) and Bianhong 01 [25] (51.07% of the genome), suggesting that the difference in genome size could be because these were missing in the cultivar assemblies. Among the classified repeats, LTR retrotransposons under class I transposable elements were the most abundant type of repetitive elements, occupying 97.3 Mb (20.34% of the genome). In contrast, DNA transposons under class II transposable elements only occupied 2.24% of the genome. These included hobo-Activator (0.62% of the genome) and mutator-like elements (0.40% of the genome). Among the transposable elements, Copia occupied the largest proportion of the genome (48.0 Mb, 10.03%), followed by Gypsy (25.3 Mb, 5.28%). Long interspersed elements occupied 0.40% of the genome, and no short interspersed nuclear element was identified in the genome.

To predict protein-coding genes, we generated a total of 48.9 Gb Illumina RNA-seq reads and 3.81 Gb Nanopore direct RNA sequencing reads from leaf, root, and stem tissues. Using both short- and long-read transcriptomic data as transcript evidence, we predicted a total of 29,004 gene models with 32,756 protein-coding transcripts in the genome of four-seeded wild lablab (Table 1). The assembly of cv. Highworth showed a similar number of genes (30,922), but a much larger number of transcripts (79,512). Whereas Bianhong 01 had a smaller number of genes (26,180) with 50,208 transcripts. Among the identified transcripts, 31,675 transcripts (91.9%) encode proteins that have predicted functional domains. For the non-coding RNA, 545 tRNAs were identified, encoding the standard 20 types of amino acid, and 1,698 lncRNA were annotated.

### Gene family analysis

A total of 26,799 protein sequences from the protein-coding genes of four-seeded wild lablab were compared with those of four other legumes, including *Medicago truncatula* (barrel medic), *Macrotyloma geocarpum* (Kersting’s groundnut), *P. vulgaris* (common bean), and *V. angularis* (adzuki bean), using *Arabidopsis thaliana* as the outgroup. In total, 24,828 lablab genes (92.6%) were assigned to 17,324 gene families, whereas 1,971 genes remained unassigned. Among the gene families of the five legume species, 13,677 were shared by all legumes, whereas 312 families with 1,090 genes were unique to lablab (Fig. 2a). Gene Ontology (GO) enrichment was performed to inspect the functions of lablab-specific genes, yielding 27 significantly (adjusted *p* < 0 .05) enriched GO terms. The enriched terms mainly included metabolomic activities, particularly the molecular functions and processes related to nitrogen fixation in legumes, such as ammonium transmembrane transporter, arginine catabolism and spermidine biosynthetic process (Additional File 2: Table S1).

**Fig. 2.**
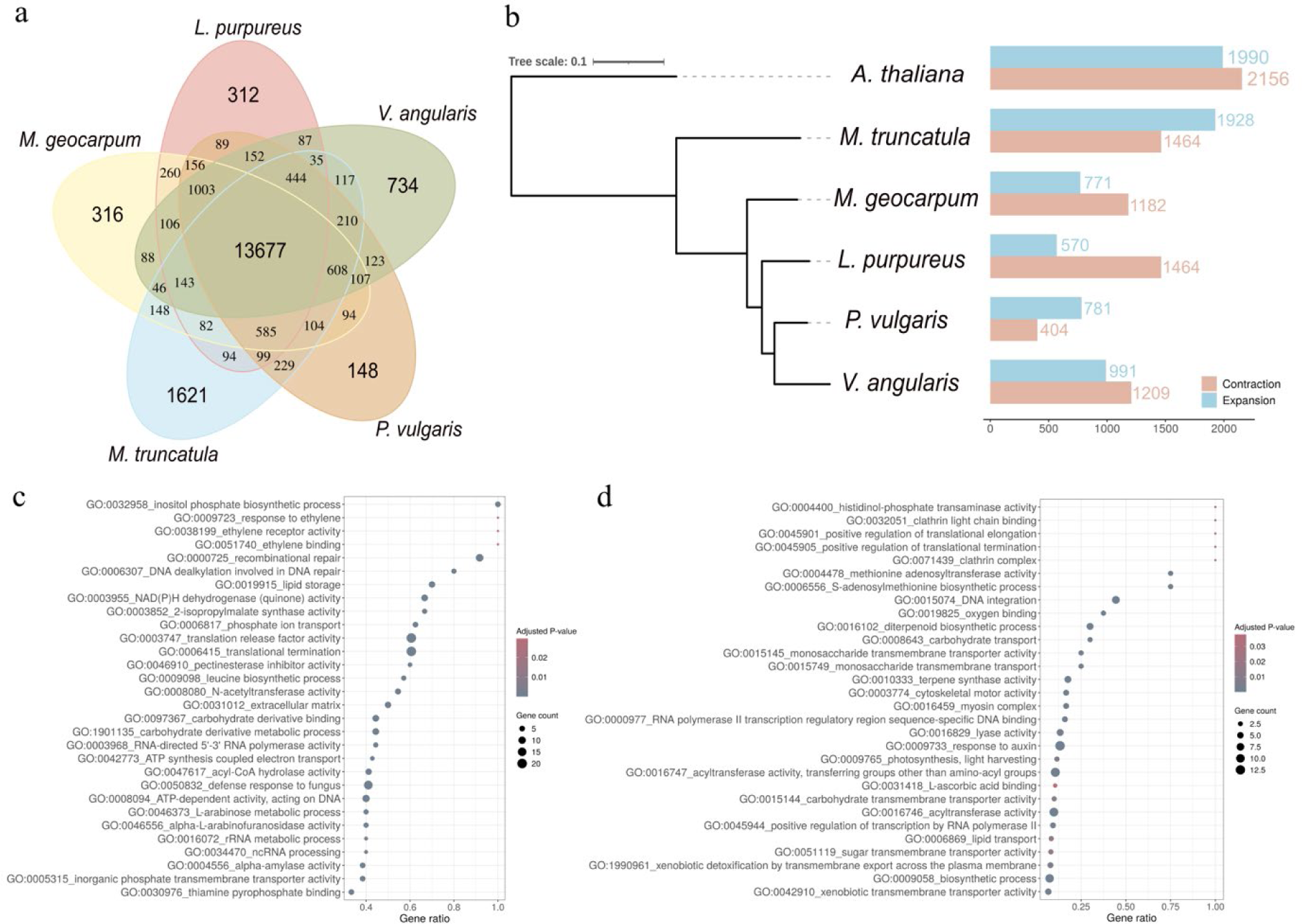
Gene family analysis of *Lablab purpureus*. (a) The shared and unique gene families in lablab compared with the four selected legume species (*Medicago truncatula*, *Macrotyloma geocarpum*, *Vigna angularis*, and *Phaseolus vulgaris*). (b) Phylogeny of lablab and the selected species showing the expanded and contracted gene families in each species. The phylogenetic tree was constructed using iTol [27]. Gene ontology terms (adjusted *p* < 0.05) enriched in the set of (c) significantly expanded and (d) contracted gene families in lablab using the one-sided over-representation analysis for gene set enrichment. The *p*-value was computed using the Fisher’s exact test, and the adjusted *p*-value was calculated using the Benjamini–Hochberg method.

By overlaying gene families on the phylogenetic relationships of lablab with the four selected legumes and *Arabidopsis*, we identified 570 expanded and 1,464 contracted gene families in lablab (Fig. 2b). Of these, 108 gene families containing 610 genes were significantly expanded (*p* < 0.05), and 376 gene families containing 281 genes were significantly contracted. Notably, 219 gene families were absent from lablab. We extracted the genes in the significantly expanded and contracted gene families and performed GO enrichment analysis. This yielded 87 and 43 significantly enriched GO terms (adjusted *p* < 0.05) for the expanded and contracted gene families, respectively. The GO terms in the expanded gene families were enriched in a diverse range of biological processes and molecular functions related to plant development and stress responses, such as response to ethylene and defense response to fungus (Fig. 2c), whereas those enriched in the contracted gene families were mainly related to cellular metabolism, transport, and regulatory functions (Fig. 2d).

### Resequencing, phylogeny, and genetic diversity

Information on the 103 lablab accessions is available in Additional File 2: Table S2. After QC there remained between 17.9 million and 43.2 million read pairs per sample (average 27.9 ± 0.8 (SE) million) except for one sample (cv. Highworth) where 50 million pre-trimmed reads from Njaci *et al*. [8] were used. The percentage of reads mapping and percentage of reads properly mapping were >95% and >84% respectively with the exception of the outgroup where these values were 57.5% and 44.7% respectively. Sequencing depth was at least 6.3X across lablab samples (3.3X for the outgroup) and average sequencing depth was 10.5X ± 0.3 (SE). After aligning reads to the newly generated wild lablab genome, 37.6 million variants (35.5 million SNPs and 2.1 million indels) were identified. After removing sites with >10% missing calls, those with MAF < 0.05 and then removing sited in LD, 161,064 sites were retained.

The Neighbor Joining (NJ) tree of all 103 samples (161,064 SNPs; Additional File 1: Fig. S1) demonstrated that of the six pairs of accessions where members of each pair was from different genebanks (see methods), four pairs appeared identical being on sister branches and showing almost zero branch length, therefore one from each pair was removed. For the other two pairs the samples were not sisters (but were very closely related) therefore both members of these were retained. The population genomic analysis therefore included 99 samples which comprised six two-seeded wild, five two-seeded domesticated, 13 four-seeded wild, and 70 four-seeded domesticated samples, plus four potentially feral samples and an outgroup.

The NJ tree (based on 160,037 SNPs) (Fig. 3a) backs up previous work confirming that the two- and four-seeded gene pools are divergent, and that wild and domesticated samples are found in each of these gene pools. Using both the NJ and STRUCTURE analyses of the four-seeded samples (Fig. 3), wild and domesticated samples are clearly divergent, with the feral samples intermediate. STRUCTURE found strong support for 2 or 7 clusters (Fig. 3b). Based on K=2 (Fig. 3c) all wild samples were grouped into one cluster and all domesticated samples to the other cluster (with >96% membership), and feral samples comprised at least 20% membership from both clusters. Based on K=7 (Fig. 3d), the wild samples form one cluster (cluster 1 in Fig. 3d) and then there are four main clusters in the domesticated group (clusters 2, 4, 6, 7). Clusters 3 and 5 represent only minor contributors to any accessions (and were never the main contributor to any individual). Cluster 2 contains only accessions from Africa, cluster 4 accessions are mainly from Africa plus two from India, cluster 6 accessions are mostly from Africa with one from Thailand and one from Israel, and accessions in cluster 7 are from around the world. It is worth noting that non-African accessions (see dots in Fig. 3a) are always nested in groups of African samples suggesting they are derived (often independently) from African domesticated samples.

**Fig. 3.**
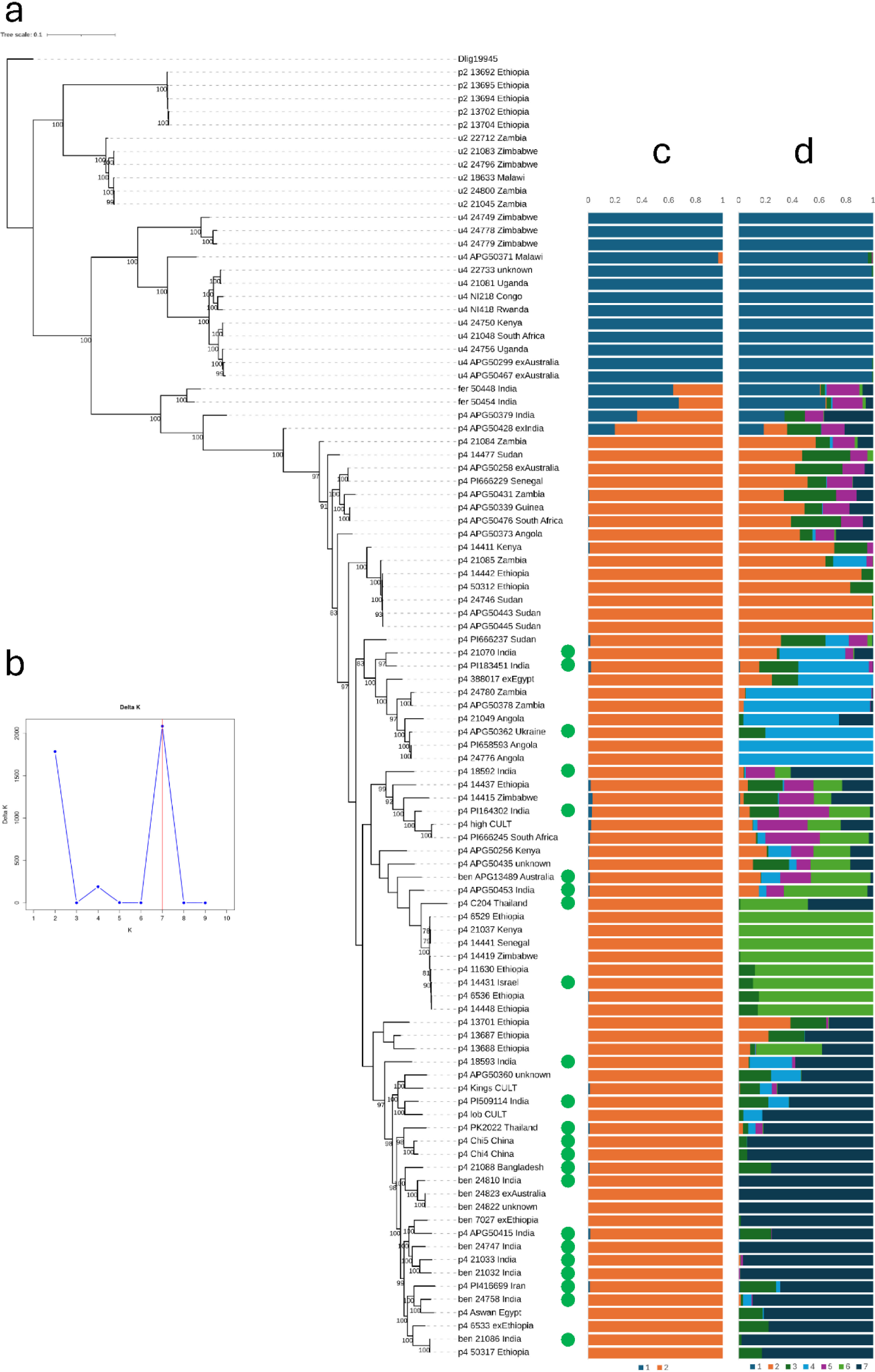
Phylogenetic and population genomic analysis. Samples are labelled as u2, u4, p2, p4 corresponding to wild (subsp. *uncinatus*) two-seeded, wild four-seeded, domesticated (subsp. *purpureus*) two-seeded and domesticated four-seeded, respectively. The NJ tree (a) was computed with 160,037 SNPs (see Methods for details). Branch support on the NJ tree is based on 100 bootstraps, and the tree is rooted on *Dipogon*. Non-African samples nested in clades of African samples are indicated with green dots. (b) shows the results of the Evanno method for inferring the number of genetic clusters (K), demonstrating that K=2 (panel c) and K=7 (panel d) fit the data better than other K. The STRUCTURE results of the four-seeded accessions (based on 54,534 SNPs) demonstrate proportional membership of each individual to one of the two (c) or seven (d) genetic clusters.

Two sets of SNP markers were identified that could differentiate all 83 four-seeded samples (Additional File 2: Table S3). Each set comprised 11 or 12 markers and could be converted to SNP assay to be used in germplasm testing in the future.

### Selective sweep

To study the domestication events in the four-seeded gene pool, we retained 6,513,958 single-nucleotide polymorphisms (SNPs) from 70 four-seeded domesticated accessions after excluding SNPs missing in more than eight samples and a MAF < 0.05. The PCA using these SNPs shows clear divergence between wild and domesticated accession (Fig. 4a, b). To identify putative selective sweeps, a genome-wide scan was carried out using a 50-kb window, dividing the genome into 9,217 regions. Using a likelihood-based strategy, we identified 397 (1,980 genes) and 79 (412 genes) windows with significant (alpha < 0.01) and strong selective sweeps within the top 5% and 1% of likelihood ratio (LR) thresholds, respectively (Fig. 4c). Several windows with evidence for a selective sweep were found to cluster consecutively and form large genomic sweep regions, mainly located on chromosomes 1, 3, 4 and 9. After merging neighbouring significant regions, this represented 313 and 43 selective sweeps in the top 5% and 1% of the LR.

**Fig. 4.**
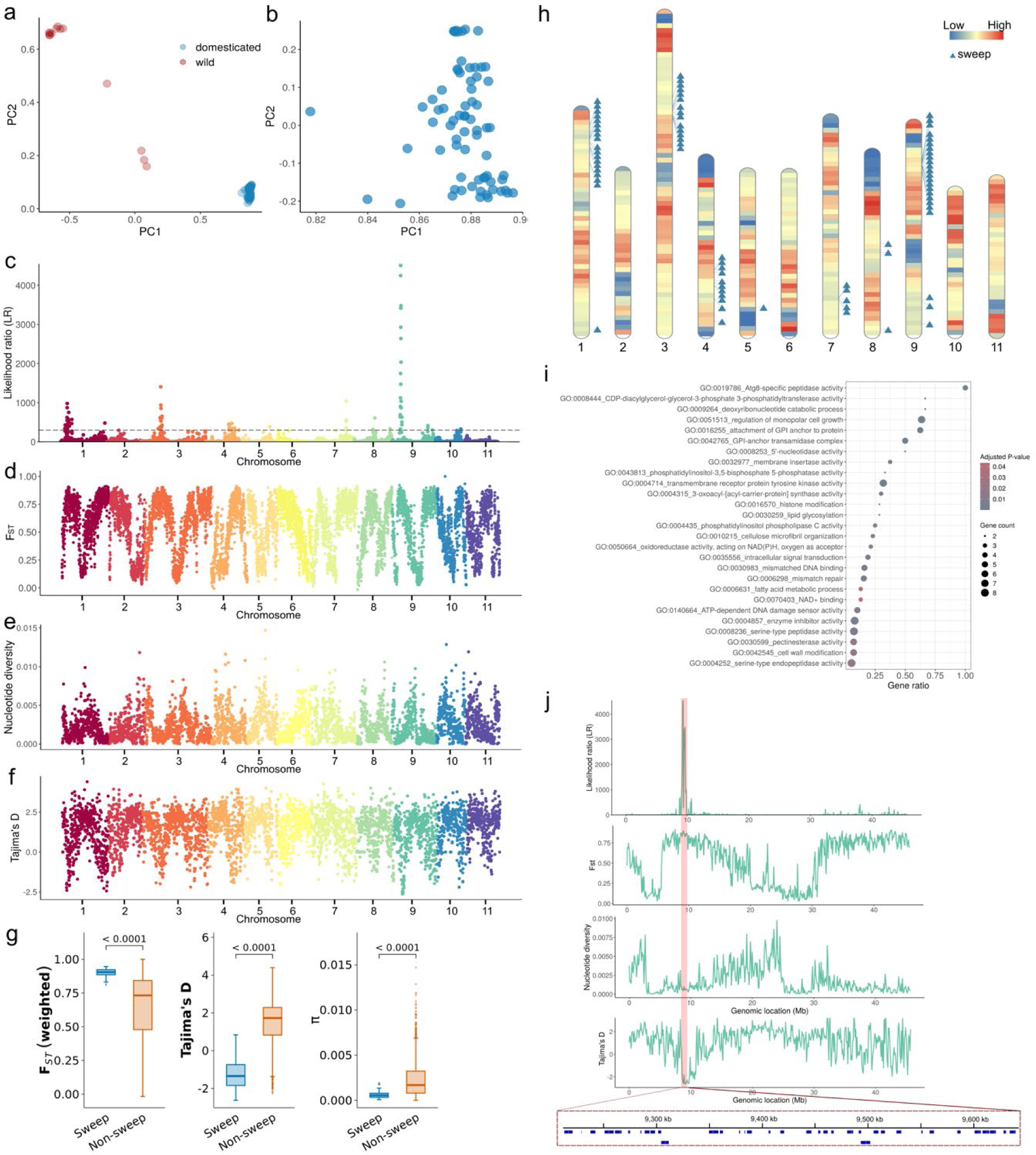
Selection of four-seeded lablab accessions. (a) Principal component analysis (PCA) of the 83 resequenced four-seeded accessions based on SNPs. (b) PCA of the 70 domesticated four-seeded lablab accessions. (c) Selective sweep analysis of the domesticated accessions. The dashed line indicates the threshold for significant sweep detection set at 1% of the highest likelihood ratio. (d) F_ST_ across 11 chromosomes between wild and domesticated populations calculated in a 100-kb sliding window with a 50-kb step. (e) Nucleotide diversity across 11 chromosomes in the domesticated population calculated in 100-kb sliding windows. (f) Tajima’s D calculated in 100-kb sliding windows. (g) Distribution of F_ST_, Tajima’s D and nucleotide diversity (π) in the strong selective sweep regions and non-sweep genomic regions. T-test and Wilcoxon rank-sum (Mann-Whitney U) test were performed. The plot was generated using Tidyplots [28]. (h) SNP density in 1-Mb windows. The triangles indicate the genomic locations of selective sweep regions within the top 1% likelihood ratio threshold. (i) GO terms (adjusted *p* < 0.05) enriched in the set of genes under selective sweep regions within the top 1% likelihood ratio threshold from the domesticated accessions. (j) Zoom-in on the selective sweep analysis of chromosome 9; the parameters from top to bottom are the likelihood ratio, F_ST_, nucleotide diversity, and Tajima’s D. The red-labeled area represents the five genomic regions exhibiting the highest likelihood ratio.

The regions with putative selective sweeps were further validated by several genome-wide population statistics, namely population divergence (F_ST_), nucleotide diversity, and Tajima’s D (TD). An average F_ST_ of 0.57 was reported between wild (13 accessions) and domesticated (70 accessions) four-seeded populations, suggesting that the two populations are highly divergent (Fig. 4d). The putative sweep regions were characterised by low nucleotide diversity and a negative Tajima’s D, suggesting potential fixation of specific domestication-related alleles and supporting the occurrence of positive selection among those regions, respectively (Fig. 4e, f). The genomic regions demonstrating putative selective sweeps had significantly lower genetic diversity, significantly lower Tajima’s D and significantly greater wild-domesticated differentiation (F_ST_) than the remainder of the genome (Fig. 4g), which would be anticipated for regions selected during domestication.

To investigate the putative positively selected genes, we focussed on the 43 selective sweeps that comprised the 79 genomic windows in the top 1% of the LR threshold. These sweeps were located in SNP-dense regions, with large sweeps on chromosomes 1, 3, 4 and 9 (Fig. 4h). The 412 genes encoding 488 transcript isoforms (Additional File 2: Table S4) in these sweep regions were subjected to GO enrichment analysis, which yielded 27 significantly enriched GO terms (adjusted *p* < 0.05). This revealed that these genes were associated with a wide range of functions, such as peptidase activity, cell wall modification, cell growth, and DNA repair mechanisms (Fig. 4i). We specifically investigated the selective sweep on chromosome 9 with the highest LR (>3,000; five 50-kb genomic regions; Fig. 4j). We identified 24 genes in this region, of which 22 have predicted functions, and 18 are potentially linked to agronomic traits, especially those related to plant growth and stress resistance (Table 2). Among 412 genes under the 43 strong selective sweep events, 36 putative pseudogenes were identified, classifying into 3 processed pseudogenes, 30 duplicated (unprocessed) pseudogenes and 3 fragmented pseudogenes. Within these putative pseudogenes, 18 genes have annotated functions according to the UniProt database (Table 3). Notably, several pseudogenes were involved in key biological processes associated to growth and stress response, including early flowering domain-containing protein, wall-associated receptor kinase and subtilisin-like protease.

**Table 2.** Candidates of agronomically related genes under positive selection in five genomic regions with the highest likelihood ratio.

| **Gene** | **Location (chr: position in bp)** | **Gene description** | **Potential implications for agronomic traits** |
| --- | --- | --- | --- |
| **Lpur_022197** | 9: 9,213,341-9,221,247 | Sucrose synthase | Fruit and seed development, response to abiotic stresses such as drought, cold, and waterlogging[29] |
| **Lpur_022198** | 9: 9,229,519-9,229,875 | Auxin responsive protein | Flower and seed development, response to stresses such as drought, high salinity, and pathogen exposure[30] |
| **Lpur_022199** | 9: 9,238,072-9,242,746 | Auxin responsive protein |  |
| **Lpur_022200** | 9: 9,251,250-9,259,982 | Respiratory burst oxidase protein | Growth and metabolism, responses to biotic and abiotic stresses such as cold stress and pathogen exposure[31, 32] |
| **Lpur_022201** | 9: 9,260,992-9,267,292 | Flavin-containing monooxygenase | Biosynthesis of auxin for plant growth and defense against pathogens[33] |
| **Lpur_022202** | 9: 9,273,877-9,275,384 | Pectinesterase inhibitor | Maintenance of cell wall integrity to enhance resistance to pathogens[34, 35] |
| **Lpur_022204** | 9: 9,289,401-9,294,012 | Pentatricopeptide repeat-containing protein | Stress response and chloroplast and pollen development[36] |
| **Lpur_022205** | 9: 9,294,630-9,299,066 | C-type lectin receptor-like tyrosine-protein kinase | Tolerance to abiotic stresses such as high salinity, drought, osmotic stress, and abscisic acid[37] |
| **Lpur_022207** | 9: 9,304,740-9,312,075 | RNA helicase | Response to abiotic and biotic stresses such as cold, heat, drought, and high salinity[38] |
| **Lpur_022219** | 9: 9,493,542-9,502,478 | von Willebrand factor type A domain-containing protein | Tolerance to biotic and abiotic stresses such as drought and high salinity[39, 40] |
| **Lpur_022220** | 9: 9,505,516-9,510,246 | PC-esterase | Resistance to high salinity, drought, and cold[41] |
| **Lpur_022222** | 9: 9,550,653-9,551,701 | LURP-one-related protein | Response to salinity, ABA stress, and pathogen[42, 43] |
| **Lpur_022223** | 9: 9,558,378-9,562,340 | Subtilisin-like protease | Response to biotic and abiotic stress such as pathogen[44], drought[45], and high salinity[46] |
| **Lpur_022224** | 9: 9,584,558-9,587,621 | Serine/threonine-protein kinase | Regulation of oil content in seeds[47], regulator of plant immune response to pathogens[48] |
| **Lpur_022225** | 9: 9,601,082-9,606,701 | Cysteine proteinase | Plant growth, regulation of protein storage in seeds, response to biotic and abiotic stresses[49] |
| **Lpur_022227** | 9: 9,615,263-9,621,047 | Tubulin-tyrosine ligase family | Regulation of cell plate and plant growth[50] |
| **Lpur_022228** | 9: 9,628,195-9,630,617 | Acyl-CoA N-acyltransferases | Regulation of meristem development, promotion of reproductive growth[51] |
| **Lpur_029109** | 9: 9,493,542-9,502,478 | Methylthioribose-1-phosphate<br>isomerase | Seed globulin plasticity in *Medicago truncatula*[52] |

**Table 3.** Putative pseudogenes with known functions under positive selection in 43 strong selective sweep events.

| **Gene** | **Transcript ID** | **Type of**<br>**pseudogene** | **UniProt ID** | **Gene description** |
| --- | --- | --- | --- | --- |
| **Lpur_007099** | Lpur_007099-T1 | Processed | A0AAN9QTK6 | Trehalose-6-phosphate synthase |
| **Lpur_000549** | Lpur_000549-T2 | Duplicated | A0A1S3TRL8 | Transcription factor MYB16 |
| **Lpur_021459** | Lpur_021459-T1 | Duplicated | A0AAQ3MDY6 | Kinesin motor domain-containing protein (Fragment) |
| **Lpur_022239** | Lpur_022239-T1 | Fragmented | V7AYP2 | Protein EARLY FLOWERING 4 domain-containing protein |
| **Lpur_011244** | Lpur_011244-T1 | Duplicated | V7C1G6 | Protein kinase domain-containing protein |
| **Lpur_011243** | Lpur_011243-T1 | Duplicated | A0A444X7Z1 | Band 7 domain-containing protein |
| **Lpur_000551** | Lpur_000551-T1 | Duplicated | V7B1T3 | Pentatricopeptide repeat-containing protein |
| **Lpur_000550** | Lpur_000550-T1 | Duplicated | V7B0U7 | NAC domain-containing protein |
| **Lpur_000638** | Lpur_000638-T1 | Duplicated | A0A0S3RSN7 | Subtilisin-like protease fibronectin type-III domain-containing protein |
| **Lpur_018543** | Lpur_018543-T1 | Duplicated | A0AAN9R339 | Carboxypeptidase |
| **Lpur_028838** | Lpur_028838-T1 | Duplicated | A0A4D6LV80 | Plasma membrane ATPase |
| **Lpur_022200** | Lpur_022200-T4 | Duplicated | A0AAQ3S7J4 | Respiratory burst oxidase-like protein E |
| **Lpur_011567** | Lpur_011567-T1 | Duplicated | A0A4D6LUG1 | tRNA (Ile)-lysidine synthetase |
| **Lpur_007158** | Lpur_007158-T2 | Fragmented | A0A4D6KU96 | Myb proto-oncogene protein |
| **Lpur_010976** | Lpur_010976-T1 | Duplicated | A0A1S3UAI1 | Pectate lyase |
| **Lpur_007152** | Lpur_007152-T1 | Duplicated | V7CEW3 | Alliinase C-terminal domain-containing protein |
| **Lpur_007151** | Lpur_007151-T1 | Duplicated | V7BFZ5 | phosphoenolpyruvate carboxylase |
| **Lpur_010962** | Lpur_010962-T1 | Processed | V7BRA7 | Wall-associated receptor kinase, galacturonan-binding domain-containing protein |

## Discussion

Underutilized crops often possess significant potential to withstand various abiotic stresses, enabling them to survive and thrive in diverse and extreme environmental conditions (i.e., geographical locations); consequently, they represent promising alternatives for sustainable agriculture in the face of global climate challenges [53]. In recent years, research on underutilized crops has steadily increased, reflecting a growing recognition of their value in sustainable food systems and the expanding scientific engagement with these crops within the academic community [54, 55]. Lablab is one such crop that is rich in nutrients, such as protein, carbohydrate, minerals, vitamins, and antioxidants, and possesses a low level of anti-nutrient factors [56, 57]. Recent studies have highlighted its potential for drought resilience [58–61] and use in pharmaceuticals [62] for treating various diseases, such as cancer [63], influenza, and COVID-19 [63], positioning lablab as a future crop for sustainable agriculture.

Crop wild relatives exhibit valuable genetic diversity that was lost during domestication. They serve as important reservoirs of key agronomic traits for crop improvement to develop cultivars with enhanced characteristics, including resilience to biotic and abiotic stresses, disease resistance, and improved nutritional quality, which are vital for the adaptation of cultivars to future climate conditions [64]. The genomes of these wild relatives play a crucial role in revealing the genetic content and identifying agronomically associated genes. Recent advances in genomic tools have made it efficient and accessible to assemble the genomes of underutilized crop and non-model organisms [21].

In this study, we generated a high-quality genome for wild lablab and conducted population genomic analyses, phylogenetics and selective sweep analysis. We applied a hybrid scaffolding approach to construct the chromosome-level assembly. When using Omni-C alone, the scaffolded lablab assembly was highly fragmented (1,344 scaffolds) with a reduced scaffold N50. After incorporating optical genome maps followed by Omni-C, 81.1% (306.4 Mb) of the scaffolds were anchored to 11 pseudomolecules, which were further subjected to fine adjustment using a homology-based strategy, constructing 96.4% (460.8 Mb) of the scaffolds into 11 pseudochromosomes. Therefore, using a combination of OGM and Omni-C for scaffolding resulted in a highly contiguous assembly for four-seeded wild lablab.

Thus far, three chromosome-level genomes have been constructed for domesticated lablab, namely the cultivar Highworth [8], a Thai landrace PK2022T020 [24] and a Chinese cultivar Bianhong 01 [25]. The assembly length of the published lablab genomes ranges from 366.4 to 463.3 Mb, whereas the wild genome generated in this study is 478.2 Mb long. The four-seeded wild genome is approximately 15 Mb, 60 Mb and 112 Mb larger than the Chinese cultivar, Highworth and Thai landrace genomes, respectively. The synteny analysis suggests a conserved genome structure and gene blocks among Highworth, Chinese cultivar and our genomes. Genome size variation can occur within species and genera [65, 66], possibly driven by differences in repetitive element content among genomes [67, 68], for example in maize [69] and the *Oryza* genus [70]. The repetitive elements occupied 143.8 Mb, 181.2 Mb, 236.6 Mb and 268.3 Mb in Thai landrace, Highworth, Chinese cultivar and four-seeded wild lablab genome respectively. This almost 2-fold difference among the repetitive elements corresponds to the observed difference in the overall genome size, indicating that the repetitive elements are the primary source of genome size variation in these lablab genomes. However, the choice of sequencing technologies and assemblers in constructing a contig-level assembly could lead to variations in the assembly length of a plant genome [71], therefore, both biological and technical factors could be playing a role. More than half of the genome (56.11%) comprised repetitive elements, whereas 51.07%, 43.36% and 39.24% of the three cultivar genomes, Bianhong 01, Highworth and PK2022T020, respectively, comprised repetitive elements. Similarly, copia elements were the most abundant type of repeats in all of the lablab genomes; in contrast, gypsy elements occupy a larger proportion than copia elements in other legume species that are closely related to lablab, including common bean [72], adzuki bean [72], and Kersting’s groundnut [73]. Despite the genome size variation, the three lablab genomes had similar numbers of genes. The four-seeded wild lablab genome contained 29,004 gene models, which is comparable to the landrace cultivar (28,511), slightly less than Highworth (30,922), but considerably higher than the Chinese cultivar (26,180).

Two independent domestication events from divergent wild populations have been previously confirmed in lablab [8] and our population genomics and phylogenetic analyses fully support this, additionally confirming that feral samples (with an intermediate wild-domesticated morphology) are indeed genetically intermediate. Our analysis shows for the first time that domesticated samples from outside Africa are always derived from African accessions. This supports that domestication took place in Africa prior to domesticated samples being transported to Southeast Asia, and that this transport occurred several times with genetically differentiated populations. Further, on a few occasions there appears to be back transport to Africa. For example, cluster 7 (blue cluster at the bottom of Fig. 3d) is primarily a Southeast Asian group of accessions (possibly derived from Ethiopian accessions), however samples from Egypt and Ethiopia are nested within this.

Here, we investigated selective sweeps and identified putative positively selected genes to unravel the domestication events in four-seeded lablab. Our analyses revealed 79 windows with evidence for a strong selective sweep (which merged into 43 selective sweeps) in the four-seeded domesticated accessions, identifying the targets of positive selection during domestication. Notably, the peak of the sweep on chromosome 9 with the greatest likelihood ratio encompassed 24 candidate genes, most of which were functionally annotated to yield-related traits and resistance to biotic and abiotic stresses. The agronomic properties of these genes have been validated in other crop species, for example, respiratory burst oxidase protein for biotic and abiotic stress tolerance in rice [31], LURP-one-related protein for response to salinity stress in *Brassica napus* [42], and subtilisin-like protease for response to drought stress in common bean [45]. Therefore, these putative positively selected genes can be candidates for breeding crops with improved productivity and stress tolerance.

Pseudogenes are non-functional genetic elements created by retrotransposition or duplication, inferring the genetic evolution history in plant species [74]. A high abundance of pseudogenes within a gene family may indicate a low selective pressure on gene whose functions are no longer vital for the species [75]. Resurrection of pseudogene could restore the function of pseudogene and create novel phenotypes [76], potentially assisting in crop breeding. A natural resurrection has been reported in plant, in which a mutation in the pseudogene of a MYB transcription factor restored the purple colour of flowers in *Petunia secreta* during speciation [77]. Therefore, among the genes under strong positive selection in lablab, we also identified putative pseudogenes. Thirty-six genes (approximately 10%) were predicted as pseudogenes in the wild reference genome and some showed similarity to pseudogenes associated with early flowering and stress response. For example, a NAC domain-containing protein mediates drought stress tolerance in legume crops [78] and wall-associated receptor kinase regulates response to salt stress in tomato [79]. The predominance of duplicated (unprocessed) pseudogenes over retroposed (processed) pseudogenes in the positively selected genes aligns with patterns observed across plant genomes in previous study [75], contrasting with mammalian genomes where retroposed pseudogenes dominate [80]. This suggests that DNA duplication events instead of retrotransposition are the major driver of pseudogenization in the lineage containing four-seeded lablab. While it is likely that each of the sweeps is only representing the population genomic effect of selection of one or two genes in the region, and not all genes, if selection was occurring for some of these pseudogenes it would indicate that the wild genome is a reservoir of latent genetic diversity that may contribute to future adaptation, demonstrating the importance of a wild reference genome in evolutionary study.

These findings collectively underscore how domestication has shaped the characteristics of stress resilience in four-seeded lablab, providing a roadmap for marker-assisted breeding to enhance stress tolerance in lablab and other legumes. Developing lablab cultivars with enhanced biotic and abiotic stress tolerance remains a top priority in breeding programs under current and future climate conditions. The first wild genomic resource of lablab, together with a population study and a set of core SNPs to differentiate the domesticated samples, enhances our understanding of genetic diversity, phylogeny, and evolutionary and domestication processes, thereby unlocking the potential of this neglected crop. Our results provide a foundation for genomic-assisted breeding of this underutilized crop aimed at improving stress resilience and crop productivity and thereby addressing the urgent need for climate-resilient crop varieties.

### Conclusions

This study presents the first high-quality reference genome with a comprehensive gene annotation for four-seeded wild lablab, revealing a complex domestication history from Africa to Asia through a population genomic analysis of 103 wild and domesticated accessions. Selective sweeps identified key genes and pseudogenes for yield and stress tolerance, providing direct genetic targets for crop breeding to accelerate crop improvement of underutilized legumes and highlighting the untapped value of wild gene pool.

## Methods

### Sample preparation and genome sequencing

The plants for WGS (ILRI accession 24779) were grown in a 1:1 mix of vermiculite and soil at the Chinese University of Hong Kong. Young leaves were collected, immediately frozen in liquid nitrogen, and stored at −80°C. High-molecular-weight (HMW) DNA was extracted from a single plant using a Nanobind plant nuclei kit (Pacific BioSciences, Menlo Park, CA, USA, Cat No. 102-302-000). Briefly, 2 g of frozen leaf tissue was homogenized in liquid nitrogen and lysed to purify the nuclei. HMW DNA was then extracted from the purified plant nuclei using a Nanobind disk. The quantity and quality of the extracted DNA were assessed using a Qubit 4 Fluorometer (Thermo Fisher Scientific, Waltham, MA, USA) and NanoDrop spectrophotometer (Thermo Fisher Scientific), respectively. Qualified DNA samples were then subjected to library preparation and PacBio HiFi sequencing on the Sequel II platform at Novogene Co., Ltd. (Beijing, China).

An Omni-C library was constructed using a Dovetail® Omni-C® Kit (Cantata Bio, Scotts Valley, CA, USA, Cat No. 21005) according to the manufacturer’s protocol. Briefly, 0.3 g of frozen leaf tissue was homogenized in liquid nitrogen and then subjected to crosslinking with 37% formaldehyde in 1× phosphate-buffered saline and nuclease digestion using endonuclease DNase I. The fragment size distribution and quantity of lysate were assessed using TapeStation D5000 HS ScreenTape (Agilent Technologies, Santa Clara, CA, USA, Cat No. 5067-5592) and a Qubit 4 Fluorometer (Thermo Fisher Scientific), respectively. The lysate that met the criteria outlined in the library preparation guidelines (fragment distribution) was used to prepare the Omni-C library. The constructed library was sequenced on Illumina NovaSeq 6000 platform (150-bp paired-end sequencing) at Novogene Co., Ltd. (Beijing, China).

Bionano OGM was performed for scaffolding. Young leaves were collected from 7-day-old seedlings growing in vermiculite at the Chinese University of Hong Kong. Ultra-high molecular weight (UHMW) DNA was extracted from 0.5 g fresh leaves using a Bionano Prep Plant Tissue DNA Isolation Kit (Bionano Genomics, San Diego, CA, USA, Cat No. 80003) following the High Polysaccharides Plant Tissue DNA Isolation Protocol (Bionano Document No. 30128, Rev C). Subsequently, 750 ng of the extracted UHMW DNA was fluorescently labelled using the Direct Label and Stain (DLS) Kit (Bionano Genomics, Cat No. 80005) following the Bionano Prep DLS Protocol (Bionano Document No. 30206, Rev G). Briefly, the UHMW DNA molecules were labeled using the Direct Label Enzyme (DLE-1) with DL-green fluorophore to generate sequence-specific patterns, followed by overnight fluorescent staining of DNA backbones to visualize the full length of the DNA molecules. The labelled UHMW DNA was loaded onto a Saphyr Chip G1.2 (Bionano Genomics, Cat No. 20319) and run on a Bionano Saphyr system (Bionano Genomics) for 36 h.

### Transcriptome sequencing

To annotate the genes of the genome, total RNA was extracted from multiple fresh tissues, including leaf, stem, and root tissues, from the same plant used for the genome sequencing for both short-read Illumina sequencing and Nanopore direct RNA sequencing. Briefly, total RNA was extracted by lysing 100 mg tissue with 1 mL RNAiso Plus (Takara Bio, San Jose, CA, USA, Cat No. 9108), followed by the addition of chloroform for phase separation. The aqueous phase containing RNA was then subjected to purification using an RNeasy Mini kit (Qiagen, Venlo, The Netherlands, Cat No. 74104) following the manufacturer’s protocol. The RNA sample was treated with DNase I (New England Biolabs, Ipswich, MA, USA, Cat No. M0303L) and purified using a Monarch RNA clean-up kit (New England Biolabs, Cat No. T2040L) to remove contaminating genomic DNA. The quality and quantity of the RNA samples were assessed using a NanoDrop spectrophotometer (Thermo Fisher Scientific) and Qubit 4 Fluorometer (Thermo Fisher Scientific), respectively. The purified RNA samples were then used to prepare strand-specific rRNA-depleted libraries and sequenced on Illumina NovaSeq 6000 platform (150-bp paired-end sequencing) at Novogene Co., Ltd. (Beijing, China). For the long-read transcriptome data, Nanopore direct RNA sequencing was performed. Poly(A) mRNA was enriched from 50 μg of total RNA using the NEBNext® poly(A) mRNA magnetic isolation module (New England Biolabs, Cat No. T2040L), followed by library preparation using a Nanopore direct RNA sequencing kit (Oxford Nanopore Technologies, Cat No. SQK-RNA002) according to the manufacturer’s protocol. The constructed libraries were subjected to sequencing on R9.4.1 flow cells (Oxford Nanopore Technologies, Oxford, UK, Cat No. FLO-MIN106D) in a MinION system for 72 h.

### *De novo* genome assembly and scaffolding

The genome heterozygosity was estimated by Jellyfish (v2.3.0) [81] and GenomeScope (v1.0) [82] using PacBio HiFi reads, following a Kmer-based statistical method. Initially, a *de novo* genome assembly was constructed by Hifiasm (v0.19.9-r616) [83] using PacBio HiFi reads with default parameters. To construct pseudomolecules from the contig-level assembly built using PacBio HiFi reads, the assembly was subjected to the first round of scaffolding using OGM data. The *de novo* assembly and scaffolding were conducted using Bionano Solve software (v3.8.1) (Bionano Genomics, Inc.). The non-haplotype setting without extend and split was applied, which is typical for plant OM scaffolding. The hybrid scaffolds from Bionano were then subjected to a second round of scaffolding using Omni-C (Hi-C) sequencing reads. The Omni-C reads were trimmed using Trim galore (v0.6.7) [84] to remove adapter sequences and low-quality bases based on a Phred score threshold of 20. The trimmed Omni-C reads were processed following the Juicer pipeline (v1.5.7) [85] and scaffolded using the 3D-DNA pipeline (v180922) [86]. This scaffolding process included the correction and orientation of contigs into pseudomolecules. The scaffolded assembly was visualized using JuiceBox (v1.11.08) [87]. Furthermore, a final step involving homology-based scaffolding conducted by RagTag (v.2.1.0) [88] using the published Highworth genome [8] as a reference was performed to further improve the contiguity of the pseudomolecules and patch physical gaps. Gaps were filled by RFfiller [89] using PacBio HiFi reads.

The 11 pseudomolecules of the lablab genome were compared with the genomes of Highworth [8] and Bianhong 01 [25] for ordering and assignment of chromosome names. Putative homologous chromosome regions and pairwise syntenic gene blocks among three genomes were identified using MCScan (v0.9.12) [90]. The genome assembly quality was assessed by BUSCO (v5.7.1) [91] using the embryophta_odb10 lineage, and the LAI was calculated using LTR_retriever (v3.0.1) [92].

### Genome annotation

Repetitive elements, namely transposable elements, small RNAs, and simple repeats, were identified using RepeatModeler (v2.0) [93] for the scaffolded genome assembly. In addition, LTR retrotransposons were identified using LTR_FINDER (v1.2) [94] and LTR_retriever (v3.0.1) [92] to generate an LTR library. This LTR library was combined with the *de novo* repeat library constructed using RepeatModeler and RepBase (RepeatMasker-Edition version 20181026), creating a comprehensive customized repeat library. The genome assembly was soft-masked with the repeat library by RepeatMasker (v4.1.0) [95], using Dfam 3.1 as the database.

To annotate the protein-coding gene model, *ab initio* gene prediction was performed on the soft-masked genome using a combination of short- and long-read transcriptome data. For short-read transcriptome data, reads were trimmed using Trim galore (v0.6.7) [84] to remove adapter sequences and low-quality bases based on a Phred score threshold of 30 and aligned to the genome using HISAT2 (v2.2.1) [96]. The raw Nanopore sequencing signals were processed by basecalling and adapter trimming using Guppy (v6.4.6) [97], and the pre-processed reads were aligned to the genome using Minimap2 (v2.24-r1122) [98]. The alignment files from short- and long-read data were merged using SAMtools (v1.17) [99] and then subjected to *de novo* transcriptome assembly using Trinity (v2.1.1) [100] and the construction of a preliminary gene annotation using PASA pipeline (v2.4.1) [101]. Using Trinity-assembled transcriptome and PASA-annotated gene models as evidence, *ab initio* gene prediction was performed with Funannotate (v1.8.9) [102], which integrated multiple gene predictors, including Augustus, GeneMark-ES, glimmerHMM, and snap, and passed all results to Evidence Modeler to construct consensus gene models. The untranslated regions were then updated per the predictions, and the gene models were adjusted based on the RNA-seq data and Trinity-assembled transcriptome. Additionally, tRNAs were identified using tRNAscan-SE (v.2.0.11) [103] with default parameters. Long non-coding RNAs (lncRNAs) were identified using CPC2 (v1.0.1) [104] and CNCI (v2) [105] with default parameters to predict the coding potential of the transcripts. Transcripts longer than 200 nucleotides and labeled as “noncoding” by both tools were further searched against the nonredundant protein database UniRef90 from UniProt using Diamond (v2.1.12) [106]. Transcripts with an e-value < 10^−10^ or identity > 95% were excluded from the lncRNAs. The completeness of the annotated gene models was evaluated by BUSCO (v5.7.1) [91] using the embryophta_odb10 lineage. Gene functions were annotated using InterProScan (v5.63-95.0) [107] by searching the protein sequences of each gene against multiple databases, such as Pfam, RefSeq, and PANTHER.

### Gene family analysis

The protein sequence of the longest transcript of each gene was extracted and compared with those from four other legumes, namely *V. angularis* (Vigan1.1), *P. vulgaris* (PhaVulg1_0), *Me. truncatula* (MtrunA17r5.0), and *Ma. geocarpum* [73], using Orthofinder (v2.5.5) [108] with *A. thaliana* (Araport 11) as an outgroup. CAFE5 (v5.1.0) [109] was used to identify the expanded and contracted gene families from the resultant gene families (with less than 100 copies of genes from each gene family) and a rooted, ultrametric species tree generated using Orthofinder. GO enrichment for the genes in the unique gene families and the significantly expanded and contracted gene families (*p* < 0.05) was performed using GSEApy (v1.1.0) [110].

### Resequencing, phylogeny, and genetic diversity

A geographically diverse group of accessions was procured from a range of genebanks, from previously published work [8, 73] and a sample from Aswan, Egypt, kindly shared by Alan Clapham. In total, this comprised six wild two-seeded, five domesticated two-seeded, 13 wild four-seeded and 74 domesticated four-seeded samples from throughout their respective ranges, plus four feral-like samples (intermediate wild-domesticated morphology) and an outgroup (total 103 samples; Additional File 2: Table S2). Within these were six pairs of domesticated four-seeded samples which were allegedly identical but from different genebanks which we included so that we could check if they were indeed identical. Plants were grown from seed in the greenhouse at the University of Southampton and DNA extracted using a CTAB-based extraction protocol [111]. After determining the quality and quantity were sufficient the samples were sent to Novogene (Cambridge, UK) for library prep (NEBNext® Ultra™ II FS DNA Library Prep Kit (Illumina) and NEBNext® Multiplex Oligos) and PE150 sequencing on an Illumina NovaSeq X Plus.

Raw reads were trimmed using Trimmomatic (v0.32) [112] and the settings LEADING:5 TRAILING:5 SLIDINGWINDOW:4:15 MINLEN:90 and then mapped to the wild lablab genome (pseudochromosomes only) using bwa-mem2 [113]. For each sample, the resultant sam file was converted to a bam file using SAMtools [114], sorted according to coordinate and duplicate reads marked using Picard (http://broadinstitute.github.io/picard). These were combined into one VCF file per chromosome using BCFtools mpileup [114] with quality settings -q 20 -Q 20. Variants were called per chromosome using BCFtools call and then the VCFs combined using BCFtools concat. BCFtools filter was then used to remove poor quality variants using ‘QUAL>30 & DP>30’. This was then thinned down to a maximum of 1 variant per 200bp and SNPs with missing data from >10 samples (∼10%) or with MAF < 0.05 were removed using VCFtools (v0.1.16) [115]. Sites in linkage disequilibrium (LD) were identified using PLINK (v 1.9) [116] with settings 50 5 0.5 and removed using VCFtools. The VCF file was then converted to a PHYLIP file and used in FastME [117] to generate a NJ tree with 100 bootstrap replicates. The tree was visualised in iTOL (https://itol.embl.de/) and rooted with *Dipogon lignosus*. After confirming that four of the six pairs of accessions did appear to be identical (see results), one of each of these four pairs was removed and the process above repeated (total 99 samples). This dataset was also used in STRUCTURE (v2.3.4) [118] to determine the most likely number of population clusters. The initial VCF file was thinned to include only these samples, indels removed, and then the same trimming approaches as above used. Finally, because of the large number of variants retained, further thinning to one SNP per 5kb was carried out (54,534 SNPs). STRUCTURE was run with five replicate runs per K (number of clusters) comprising 50,000 iterations after a burn-in of 20,000. The Evanno method [119] was used to infer the most likely number of genetic clusters. Results were then plotted using the online CLUMPAK server (http://clumpak.tau.ac.il/) [120].

We used CoreSNP (https://github.com/admy55/CoreSNP) [121] to identify two sets of SNPs that could differentiate all 4-seeded individuals.

### Identification of selective sweeps

Selective sweeps in the domesticated population were detected using a genome-wide scan performed using Sweepfinder2 (v1.0) [122], which calculated the LR in a 50-kb window across the genome. To further validate the genomic regions of selective sweeps, we assessed the genome-wide population statistics, including TD, genetic diversity (π), and population divergence (F_ST_), using VCFtools (v0.1.16) [115] in 100-kb sliding windows. Putative genomic regions under positive selection were selected based on the sweep intensity (α) and LR. The threshold for classifying significant selective sweeps was set at the 99^th^ percentile, whereby the regions falling within the top 1% LR threshold with α < 0.01 were considered as candidate regions of strong selective sweeps. Genes within those strong sweep regions were identified using bedtools (v2.29.1) [123]. GO enrichment for the genes in the strong selective sweep regions was performed using GSEApy (v1.1.0) [110].

Pseudogenes within the windows with the top 1% LR were identified by homology-based approach in a combination of two computational tools, namely tBLASTn (v2.14.1) [124] and PseudoPipe [125]. First, the protein sequences were used as queries in a tBLASTn search against the four-seeded wild genome, in which the regions with exon and repetitive sequences were hard-masked using bedtools (v2.29.1). Hits with e-value < 10^−10^ and identity > 60%) from tBLASTn were retained as initial candidates of pseudogene. Second, PseudoPipe were further used to validate and classify putative pseudogenes into three categories, namely processed, duplicated and fragmented pseudogenes using default parameters. Briefly, a BLAST search was performed using the protein sequences as queries against the repeat-masked genome, following by refining alignment using a more accurate program, tfasty in PseudoPipe. Hits with an e-value < 10^−10^, query coverage > 50% and identity of amino acid sequence > 30% from PseudoPipe were retained as preliminary candidates. Those candidates identified by both methods were defined as high-confidence pseudogenes.

## Declarations

### Ethics approval and consent to participate

Not applicable

### Consent for publication

Not applicable

## Data availability

The genome assembly has been deposited in NCBI GenBank under the accession JBQWFI000000000 and the Genome Warehouse in National Genomics Data Center, Beijing Institute of Genomics, Chinese Academy of Sciences/China National Center for Bioinformation, under accession GWHGPXT00000000.1. The raw PacBio, Omni-C, Illumina RNA sequencing and Nanopore direct RNA sequencing data have been deposited in the NCBI Sequence Read Archive under BioProject accession PRJNA1314385 and Genome Sequence Archive (GSA) under accession CRA029574, with a BioProject accession PRJCA045617. The genome assembly and gene annotation files have been additionally deposited and are publicly available in the CUHK data repository and figshare. Resequencing data in fastq format have been deposited in the NCBI Sequence Read Archive under BioProject accession PRJNA1344857.

## Author Contributions

M.A.C. and T.-F.C. conceived and planned the experiments. T.-Y.C., D.A.T.B., Y.Y., Y.Z. and T.-F.C. performed DNA extraction, genome sequencing and construction of genome assembly. T.-Y.C. performed genome annotation, gene family and selective sweep analyses. M.A.C. carried out resequencing and performed population and phylogenetic analyses. P.S. provided materials and expertise. T.-Y.C., D.A.T.B., M.A.C. and T.-F.C. wrote the original draft of the manuscript. All authors reviewed and approved the final manuscript.

## Acknowledgements

M.A.C. acknowledges the use of the IRIDIS High Performance Computing Facility at the University of Southampton, and associated support services. We thank all the seed banks named in Table S2 and Alan Clapham for sending seed.

## Supplementary Information

Additional file 1: Fig. S1. Phylogenetic analysis of the full 103 samples. Samples are labelled as u2, u4, p2, p4 corresponding to wild (subsp. *uncinatus*) two-seeded, wild four-seeded, domesticated (subsp. *purpureus*) two-seeded and domesticated four-seeded, respectively. The NJ tree was computed with 161,064 SNPs (see Methods for details). Branch support on the NJ tree is based on 100 bootstraps, and the tree is rooted on *Dipogon*. Six alleged pairs of samples were included to examine if samples in different seedbanks were indeed identical (see Additional File 2: Table S2).

Additional file 2: Table S1. GO enrichment of gene families unique in lablab. Table S2. Information on the 103 accessions used, their origins and seedbanks. Also given are the number of reads used (after QC), the percentage mapping, and the depth. Table S3. Two sets of core SNPs that can be used to differentiate all 4-seeded samples. Table S4. Putative positively selected genes under 43 strong selective sweeps (top 1% of the LR threshold).

## References

1. Carr TW, Mkuhlani S, Segnon AC, Ali Z, Zougmoré R, Dangour AD, Green R, Scheelbeek P. Climate change impacts and adaptation strategies for crops in West Africa: a systematic review. Environmental research letters. 2022;17:53001.

2. Zhao C, Liu B, Piao S, Wang X, Lobell DB, Huang Y, Huang M, Yao Y, Bassu S, Ciais P, et al. Temperature increase reduces global yields of major crops in four independent estimates. Proc Natl Acad Sci U S A. 2017;114:9326–9331.

3. Mustafa MA, Mayes S, Massawe F, Sarkar A, Sensarma SR, vanLoon GW. Crop Diversification Through a Wider Use of Underutilised Crops: A Strategy to Ensure Food and Nutrition Security in the Face of Climate Change. In. Cham: Springer International Publishing; 2019: 125–149

4. Chivenge P, Mabhaudhi T, Modi AT, Mafongoya P. The Potential Role of Neglected and Underutilised Crop Species as Future Crops under Water Scarce Conditions in Sub-Saharan Africa. Int J Environ Res Public Health. 2015;12:5685–5711.

5. Mayes S, Massawe FJ, Alderson PG, Roberts JA, Azam-Ali SN, Hermann M. The potential for underutilized crops to improve security of food production. J Exp Bot. 2012;63:1075–1079.

6. Maass BL, Knox MR, Venkatesha SC, Angessa TT, Ramme S, Pengelly BC. Lablab purpureus—A Crop Lost for Africa? Tropical plant biology. 2010;3:123–135.

7. Missanga JS, Venkataramana PB, Ndakidemi PA. Recent developments in Lablab purpureus genomics: A focus on drought stress tolerance and use of genomic resources to develop stress-resilient varieties. Legume science. 2021;3:n/a.

8. Njaci I, Waweru B, Kamal N, Muktar MS, Fisher D, Gundlach H, Muli C, Muthui L, Maranga M, Kiambi D, et al. Chromosome-level genome assembly and population genomic resource to accelerate orphan crop lablab breeding. Nature communications. 2023;14:1915–1915.

9. Maass BL, Robotham O, Chapman MA. Evidence for two domestication events of hyacinth bean (Lablab purpureus (L.) Sweet): a comparative analysis of population genetic data. Genetic resources and crop evolution. 2017;64:1221–1230.

10. Kongjaimun A, Takahashi Y, Yoshioka Y, Tomooka N, Mongkol R, Somta P. Molecular Analysis of Genetic Diversity and Structure of the Lablab (Lablab purpureus (L.) Sweet) Gene Pool Reveals Two Independent Routes of Domestication. Plants (Basel). 2022;12.

11. Minde JJ, Venkataramana PB, Matemu AO. Dolichos Lablab-an underutilized crop with future potentials for food and nutrition security: a review. Crit Rev Food Sci Nutr. 2021;61:2249–2261.

12. Singh A, Abhilash PC. Varietal dataset of nutritionally important Lablab purpureus (L.) Sweet from Eastern Uttar Pradesh, India. Data Brief. 2019;24:103935.

13. Hossain S, Ahmed R, Bhowmick S, Mamun AA, Hashimoto M. Proximate composition and fatty acid analysis of Lablab purpureus (L.) legume seed: implicates to both protein and essential fatty acid supplementation. Springerplus. 2016;5:1899.

14. Yao L, Xia Z, Tang P, Deng J, Hao E, Du Z, Jia F, Wang X, Li Z, Fan L, Hou X. Botany, traditional uses, phytochemistry, pharmacology, edible uses, and quality control of Lablab semen Album: A systematic review. J Ethnopharmacol. 2024;334:118507.

15. Morris JB. Morphological and reproductive characterization in hyacinth bean, Lablab purpureus (L.) sweet germplasm with clinically proven nutraceutical and pharmaceutical traits for use as a medicinal food. J Diet Suppl. 2009;6:263–279.

16. Teshome A, Habte E, Cheema J, Mekasha A, Lire H, Muktar MS, Quiroz-Chavez J, Domoney C, Jones CS. A population genomics approach to unlock the genetic potential of lablab (Lablab purpureus (L.) Sweet), an underutilized tropical forage crop. BMC Genomics. 2024;25:1241.

17. Gonal B, Sampangi R, Mugali KP, Chindi SB, Chandana BR, Satish H, Prashantha V, Karthik N, Sindhu D, Kemparaju M, Sinchana BV. Discovery and validation of SSR marker-based QTL governing fresh pod yield in dolichos bean (Lablab purpureus L. Sweet). Sci Rep. 2025;15:8613.

18. Kalpana MP, Ramesh S, Siddu CB, Basanagouda G, Madhusudan K, Sathish H, Sindhu D, Kemparaju M, Anilkumar C. Low density marker-based effectiveness and efficiency of early-generation genomic selection relative to phenotype-based selection in dolichos bean (Lablab purpureus L. Sweet). Plant Genome. 2025;18:e70039.

19. Ross-Ibarra J, Morrell PL, Gaut BS. Plant domestication, a unique opportunity to identify the genetic basis of adaptation. Proc Natl Acad Sci U S A. 2007;104 Suppl 1:8641–8648.

20. Pourkheirandish M, Golicz AA, Bhalla PL, Singh MB. Global Role of Crop Genomics in the Face of Climate Change. Frontiers in plant science. 2020;11.

21. Chapman MA, He Y, Zhou M. Beyond a reference genome: pangenomes and population genomics of underutilized and orphan crops for future food and nutrition security. The New phytologist. 2022;234:1583–1597.

22. Krishna TPA, Veeramuthu D, Maharajan T, Soosaimanickam M. The Era of Plant Breeding: Conventional Breeding to Genomics-assisted Breeding for Crop Improvement. Curr Genomics. 2023;24:24–35.

23. Chang Y, Liu H, Liu M, Liao X, Sahu SK, Fu Y, Song B, Cheng S, Kariba R, Muthemba S, et al. The draft genomes of five agriculturally important African orphan crops. Gigascience. 2019;8.

24. Pootakham W, Somta P, Kongkachana W, Naktang C, Sonthirod C, U-Thoomporn S, Yoocha T, Phadphon P, Tangphatsornruang S. A de novo chromosome-scale assembly of the Lablab purpureus genome. Frontiers in plant science. 2024;15:1347744–1347744.

25. Wang L, Jiang X, Jiao W, Mao J, Ye W, Cao Y, Chen Q, Song Q. Pangenome analysis provides insights into legume evolution and breeding. Nat Genet. 2025;57:2052–2061.

26. Ou S, Chen J, Jiang N. Assessing genome assembly quality using the LTR Assembly Index (LAI). Nucleic acids research. 2018;46:e126–e126.

27. Letunic I, Bork P. Interactive Tree of Life (iTOL) v6: recent updates to the phylogenetic tree display and annotation tool. Nucleic acids research. 2024;52:W78–W82.

28. Engler JB. Tidyplots empowers life scientists with easy code-based data visualization. Imeta. 2025;4:e70018.

29. Li J, Hu Y, Hu J, Xie Q, Chen X, Qi X. Sucrose synthase: An enzyme with multiple roles in plant physiology. J Plant Physiol. 2024;303:154352.

30. Bao D, Chang S, Li X, Qi Y. Advances in the study of auxin early response genes: Aux/IAA, GH3, and SAUR. The Crop journal. 2024;12:964–978.

31. Khaskhali S, Xiao X, Zhang Z, Solangi F, Hussain S, Chen Y. Expression profile and characterization of respiratory burst oxidase homolog genes in rice under MeJA, SA and Xoo treatments. Sci Rep. 2025;15:5936.

32. Baxter A, Mittler R, Suzuki N. ROS as key players in plant stress signalling. J Exp Bot. 2014;65:1229–1240.

33. Schlaich NL. Flavin-containing monooxygenases in plants: looking beyond detox. Trends Plant Sci. 2007;12:412–418.

34. Liu N, Sun Y, Pei Y, Zhang X, Wang P, Li X, Li F, Hou Y. A Pectin Methylesterase Inhibitor Enhances Resistance to Verticillium Wilt. Plant Physiol. 2018;176:2202–2220.

35. Lionetti V, Fabri E, De Caroli M, Hansen AR, Willats WG, Piro G, Bellincampi D. Three Pectin Methylesterase Inhibitors Protect Cell Wall Integrity for Arabidopsis Immunity to Botrytis. Plant Physiol. 2017;173:1844–1863.

36. Meng L, Du M, Zhu T, Li G, Ding Y, Zhang Q. PPR proteins in plants: roles, mechanisms, and prospects for rice research. Front Plant Sci. 2024;15:1416742.

37. Vaid N, Macovei A, Tuteja N. Knights in action: lectin receptor-like kinases in plant development and stress responses. Mol Plant. 2013;6:1405–1418.

38. Li X, Li C, Zhu J, Zhong S, Zhu H, Zhang X. Functions and mechanisms of RNA helicases in plants. J Exp Bot. 2023;74:2295–2310.

39. Karkute SG, Kumar V, Tasleem M, Mishra DC, Chaturvedi KK, Rai A, Sevanthi AM, Gaikwad K, Sharma TR, Solanke AU. Genome-Wide Analysis of von Willebrand Factor A Gene Family in Rice for Its Role in Imparting Biotic Stress Resistance with Emphasis on Rice Blast Disease. Rice science. 2022;29:375–384.

40. Wang Q, Tian S, Zhang X, Zhang Y, Wang Y, Xie S. Insights into the tolerant function of VWA proteins in terms of expression analysis and RGLG5-VWA crystal structure. Plant Physiol Biochem. 2024;214:108864.

41. Anderson AC, Stangherlin S, Pimentel KN, Weadge JT, Clarke AJ. The SGNH hydrolase family: a template for carbohydrate diversity. Glycobiology. 2022;32:826–848.

42. Yang S, Chen J, Ding Y, Huang Q, Chen G, Ulhassan Z, Wei J, Wang J. Genome-wide investigation and expression profiling of LOR gene family in rapeseed under salinity and ABA stress. Front Plant Sci. 2023;14:1197781.

43. Baig A. Role of Arabidopsis LOR1 (LURP-one related one) in basal defense against Hyaloperonospora arabidopsidis. Physiological and molecular plant pathology. 2018;103:71–77.

44. Figueiredo A, Monteiro F, Sebastiana M. Subtilisin-like proteases in plant-pathogen recognition and immune priming: a perspective. Front Plant Sci. 2014;5:739.

45. Budič M, Sabotič J, Meglič V, Kos J, Kidrič M. Characterization of two novel subtilases from common bean (Phaseolus vulgaris L.) and their responses to drought. Plant Physiol Biochem. 2013;62:79–87.

46. Liu JX, Srivastava R, Che P, Howell SH. Salt stress responses in Arabidopsis utilize a signal transduction pathway related to endoplasmic reticulum stress signaling. Plant J. 2007;51:897–909.

47. Ramachandiran I, Vijayakumar A, Ramya V, Rajasekharan R. Arabidopsis serine/threonine/tyrosine protein kinase phosphorylates oil body proteins that regulate oil content in the seeds. Sci Rep. 2018;8:1154.

48. Lin ZJ, Liebrand TW, Yadeta KA, Coaker G. PBL13 Is a Serine/Threonine Protein Kinase That Negatively Regulates Arabidopsis Immune Responses. Plant Physiol. 2015;169:2950–2962.

49. Grudkowska M, Zagdańska B. Multifunctional role of plant cysteine proteinases. Acta Biochim Pol. 2004;51:609–624.

50. Zhang K, Zhu X, Durst S, Hohenberger P, Han MJ, An G, Sahi VP, Riemann M, Nick P. A rice tubulin tyrosine ligase-like 12 protein affects the dynamic and orientation of microtubules. J Integr Plant Biol. 2021;63:848–864.

51. Walla A, Wilma van Esse G, Kirschner GK, Guo G, Brünje A, Finkemeier I, Simon R, von Korff M. An Acyl-CoA N-Acyltransferase Regulates Meristem Phase Change and Plant Architecture in Barley. Plant Physiol. 2020;183:1088–1109.

52. Cartelier K, Aimé D, Ly Vu J, Combes-Soia L, Labas V, Prosperi JM, Buitink J, Gallardo K, Le Signor C. Genetic determinants of seed protein plasticity in response to the environment in Medicago truncatula. Plant J. 2021;106:1298–1311.

53. Kaur S, Kaur G, Kumari A, Ghosh A, Singh G, Bhardwaj R, Kumar A, Riar A. Resurrecting forgotten crops: Food-based products from potential underutilized crops a path to nutritional security and diversity. Future foods : a dedicated journal for sustainability in food science. 2025;11:100585.

54. Ndlovu M, Scheelbeek P, Ngidi M, Mabhaudhi T. Underutilized crops for diverse, resilient and healthy agri-food systems: a systematic review of sub-Saharan Africa. Front Sustain Food Syst. 2024;8.

55. Adelabu DB, Franke AC. Status of underutilized crop production: Its potentials for mitigating food insecurity. Agronomy journal. 2023;115:2174–2193.

56. Pandey DK, Singh S, Dubey SK, Mehra TS, Dixit S, Sawargaonkar G. Nutrient profiling of lablab bean (Lablab purpureus) from north-eastern India: A potential legume for plant-based meat alternatives. Journal of food composition and analysis. 2023;119:105252.

57. Bhattacharya S, Malleshi NG. Physical, chemical and nutritional characteristics of premature-processed and matured green legumes. J Food Sci Technol. 2012;49:459–466.

58. Akello M, Nyaboga EN, Badji A, Rubaihayo P. Deciphering the morpho-physiological and biochemical responses in Lablab purpureus (L.), Sweet, seedlings to water stress. South African journal of botany. 2023;162:412–424.

59. Missanga JS, Venkataramana PB. Lablab (Lablab Purpureus L. Sweet): Current Developments in Breeding and Prospects for Drought Tolerance, Improved Yield, and Sustainable Production in Sub-Saharan Africa (SSA). In Marker-Assisted Breeding in Legumes for Drought Tolerance. Edited by Nadeem MA, Baloch FS, Bantis F, Fiaz S, Aasim M. Singapore: Springer Nature Singapore; 2025: 345–371

60. Tasnia N, Haque RB, Islam MM, Haque MA. Morphological and Molecular Characterizations of Country Bean (Lablab purpureus L.) Genotypes for Drought Tolerance. American Journal of Plant Sciences. 2024;15:1069–1090.

61. Kokila S, Devaraj VR. A comparative study of ESTs induced under drought and salinity stress in Hyacinth bean (Lablab purpureus). American Journal of Plant Sciences. 2021;12:840–857.

62. Zhou J, Wang W, Zhang Z, Zhu G, Qiao J, Guo S, Bai Y, Zhao C, Teng C, Qin P, et al. An underutilized bean: hyacinth bean [Lablab purpureus (L.) sweet]: bioactive compounds, functional activity, and future food prospect and applications. J Sci Food Agric. 2025;105:701–720.

63. Liu YM, Shahed-Al-Mahmud M, Chen X, Chen TH, Liao KS, Lo JM, Wu YM, Ho MC, Wu CY, Wong CH, et al. A Carbohydrate-Binding Protein from the Edible Lablab Beans Effectively Blocks the Infections of Influenza Viruses and SARS-CoV-2. Cell Rep. 2020;32:108016.

64. Bohra A, Kilian B, Sivasankar S, Caccamo M, Mba C, McCouch SR, Varshney RK. Reap the crop wild relatives for breeding future crops. Trends Biotechnol. 2022;40:412–431.

65. Huang H, Tong Y, Zhang QJ, Gao LZ. Genome size variation among and within Camellia species by using flow cytometric analysis. PLoS One. 2013;8:e64981.

66. Dai SF, Zhu XG, Hutang GR, Li JY, Tian JQ, Jiang XH, Zhang D, Gao LZ. Genome Size Variation and Evolution Driven by Transposable Elements in the Genus Oryza. Front Plant Sci. 2022;13:921937.

67. Stelzer CP, Blommaert J, Waldvogel AM, Pichler M, Hecox-Lea B, Mark Welch DB. Comparative analysis reveals within-population genome size variation in a rotifer is driven by large genomic elements with highly abundant satellite DNA repeat elements. BMC Biol. 2021;19:206.

68. Cang FA, Welles SR, Wong J, Ziaee M, Dlugosch KM. Genome size variation and evolution during invasive range expansion in an introduced plant. Evol Appl. 2024;17:e13624.

69. Wang Q, Dooner HK. Remarkable variation in maize genome structure inferred from haplotype diversity at the bz locus. Proc Natl Acad Sci U S A. 2006;103:17644–17649.

70. Long W, He Q, Wang Y, Wang Y, Wang J, Yuan Z, Wang M, Chen W, Luo L, Luo L, et al. Genome evolution and diversity of wild and cultivated rice species. Nat Commun. 2024;15:9994.

71. Murigneux V, Rai SK, Furtado A, Bruxner TJC, Tian W, Harliwong I, Wei H, Yang B, Ye Q, Anderson E, et al. Comparison of long-read methods for sequencing and assembly of a plant genome. Gigascience. 2020;9.

72. Chu L, Yang K, Chen C, Zhao B, Hou Y, Wang W, Zhao P, Wang K, Wang B, Xiao Y, et al. Chromosome-level reference genome and resequencing of 322 accessions reveal evolution, genomic imprint and key agronomic traits in adzuki bean. Plant Biotechnol J. 2024;22:2173–2185.

73. Cheung TY, Kafoutchoni KM, Agoyi EE, Chan TF, Chapman MA. Exceptionally low genomic diversity in the underutilised legume Kersting’s groundnut. Nat Commun. 2025;16:5183.

74. Xie J, Chen S, Xu W, Zhao Y, Zhang D. Origination and Function of Plant Pseudogenes. Plant Signal Behav. 2019;14:1625698.

75. Mascagni F, Usai G, Cavallini A, Porceddu A. Structural characterization and duplication modes of pseudogenes in plants. Sci Rep. 2021;11:5292.

76. Yadav S, Kalwan G, Meena S, Gill SS, Yadava YK, Gaikwad K, Jain PK. Unravelling the due importance of pseudogenes and their resurrection in plants. Plant Physiol Biochem. 2023;203:108062.

77. Esfeld K, Berardi AE, Moser M, Bossolini E, Freitas L, Kuhlemeier C. Pseudogenization and Resurrection of a Speciation Gene. Curr Biol. 2018;28:3776–3786.e3777.

78. Singh S, Kudapa H, Garg V, Varshney RK. Comprehensive analysis and identification of drought-responsive candidate NAC genes in three semi-arid tropics (SAT) legume crops. BMC Genomics. 2021;22:289.

79. Meco V, Egea I, Ortíz-Atienza A, Drevensek S, Esch E, Yuste-Lisbona FJ, Barneche F, Vriezen W, Bolarin MC, Lozano R, Flores FB. The Salt Sensitivity Induced by Disruption of Cell Wall-Associated Kinase 1 (SlWAK1) Tomato Gene Is Linked to Altered Osmotic and Metabolic Homeostasis. Int J Mol Sci. 2020;21.

80. Podlaha O, Zhang J. Processed pseudogenes: the ‘fossilized footprints’ of past gene expression. Trends Genet. 2009;25:429–434.

81. Marcais G, Kingsford C. A fast, lock-free approach for efficient parallel counting of occurrences of k-mers. Bioinformatics. 2011;27:764–770.

82. Ranallo-Benavidez TR, Jaron KS, Schatz MC. GenomeScope 2.0 and Smudgeplot for reference-free profiling of polyploid genomes. Nature communications. 2020;11:1432–1432.

83. Cheng H, Concepcion GT, Feng X, Zhang H, Li H. Haplotype-resolved de novo assembly using phased assembly graphs with hifiasm. Nature methods. 2021;18:170–175.

84. Trim Galore!: A wrapper around Cutadapt and FastQC to consistently apply adapter and quality trimming to FastQ files, with extra functionality for RRBS data [http://www.bioinformatics.babraham.ac.uk/projects/trim_galore/]

85. Durand NC, Shamim MS, Machol I, Rao SSP, Huntley MH, Lander ES, Aiden EL. Juicer Provides a One-Click System for Analyzing Loop-Resolution Hi-C Experiments. Cell systems. 2016;3:95–98.

86. Dudchenko O, Batra SS, Omer AD, Nyquist SK, Hoeger M, Durand NC, Shamim MS, Machol I, Lander ES, Aiden AP, Aiden EL. De novo assembly of the Aedes aegypti genome using Hi-C yields chromosome-length scaffolds. Science (American Association for the Advancement of Science). 2017;356:92–95.

87. Durand NC, Robinson JT, Shamim MS, Machol I, Mesirov JP, Lander ES, Aiden EL. Juicebox Provides a Visualization System for Hi-C Contact Maps with Unlimited Zoom. Cell systems. 2016;3:99–101.

88. Alonge M, Lebeigle L, Kirsche M, Jenike K, Ou S, Aganezov S, Wang X, Lippman ZB, Schatz MC, Soyk S. Automated assembly scaffolding using RagTag elevates a new tomato system for high-throughput genome editing. Genome Biology. 2022;23:258–258.

89. Midekso FD, Yi G. RFfiller: a robust and fast statistical algorithm for gap filling in draft genomes. PeerJ (San Francisco, CA). 2022;10:e14186–e14186.

90. Tang H, Wang X, Bowers JE, Ming R, Alam M, Paterson AH. Unraveling ancient hexaploidy through multiply-aligned angiosperm gene maps. Genome Research. 2008;18:1944–1954.

91. Simão FA, Waterhouse RM, Ioannidis P, Kriventseva EV, Zdobnov EM. BUSCO: assessing genome assembly and annotation completeness with single-copy orthologs. Bioinformatics. 2015;31:3210–3212.

92. Ou S, Jiang N. LTR_retriever: A Highly Accurate and Sensitive Program for Identification of Long Terminal Repeat Retrotransposons. Plant physiology (Bethesda). 2018;176:1410–1422.

93. Flynn JM, Hubley R, Goubert C, Rosenb J, Clark AG, Feschotte C, Smit AF. RepeatModeler2 for automated genomic discovery of transposable element families. Proceedings of the National Academy of Sciences - PNAS. 2020;117:9451–9457.

94. Ou S, Jiang N. LTR_FINDER_parallel: parallelization of LTR_FINDER enabling rapid identification of long terminal repeat retrotransposons. Mobile DNA. 2019;10:48–48.

95. RepeatMasker Open-4.0. [https://repeatmasker.org]

96. Kim D, Paggi JM, Park C, Bennett C, Salzberg SL. Graph-based genome alignment and genotyping with HISAT2 and HISAT-genotype. Nature biotechnology. 2019;37:907–915.

97. Wick RR, Judd LM, Holt KE. Performance of neural network basecalling tools for Oxford Nanopore sequencing. Genome Biology. 2019;20:129–129.

98. Li H. Minimap2: pairwise alignment for nucleotide sequences. Bioinformatics. 2018;34:3094–3100.

99. Danecek P, Bonfield JK, Liddle J, Marshall J, Ohan V, Pollard MO, Whitwham A, Keane T, McCarthy SA, Davies RM, Li H. Twelve years of SAMtools and BCFtools. Gigascience. 2021;10.

100. Grabherr MG, Haas BJ, Yassour M, Levin JZ, Thompson DA, Amit I, Adiconis X, Fan L, Raychowdhury R, Zeng Q, et al. Trinity: reconstructing a full-length transcriptome without a genome from RNA-Seq data. Nature biotechnology. 2011;29:644-652.

101. Haas BJ, Delcher AL, Mount SM, Wortman JR, Smith Jr RK, Hannick LI, Maiti R, Ronning CM, Rusch DB, Town CD, et al. Improving the Arabidopsis genome annotation using maximal transcript alignment assemblies. Nucleic acids research. 2003;31:5654–5666.

102. Palmer JM, Stajich J. Funannotate v1.8.1: eukaryotic genome annotation. 2020.

103. Lowe TM, Eddy SR. tRNAscan-SE: A Program for Improved Detection of Transfer RNA Genes in Genomic Sequence. Nucleic acids research. 1997;25:955–964.

104. Kang YJ, Yang DC, Kong L, Hou M, Meng YQ, Wei L, Gao G. CPC2: a fast and accurate coding potential calculator based on sequence intrinsic features. Nucleic Acids Res. 2017;45:W12–w16.

105. Sun L, Luo H, Bu D, Zhao G, Yu K, Zhang C, Liu Y, Chen R, Zhao Y. Utilizing sequence intrinsic composition to classify protein-coding and long non-coding transcripts. Nucleic Acids Res. 2013;41:e166.

106. Buchfink B, Xie C, Huson DH. Fast and sensitive protein alignment using DIAMOND. Nat Methods. 2015;12:59–60.

107. Jones P, Binns D, Chang H-Y, Fraser M, Li W, McAnulla C, McWilliam H, Maslen J, Mitchell A, Nuka G, et al. InterProScan 5: genome-scale protein function classification. Bioinformatics. 2014;30:1236–1240.

108. Emms DM, Kelly S. OrthoFinder: phylogenetic orthology inference for comparative genomics. Genome Biology. 2019;20:238–238.

109. Mendes FK, Vanderpool D, Fulton B, Hahn MW. CAFE 5 models variation in evolutionary rates among gene families. Bioinformatics. 2021;36:5516–5518.

110. Fang Z, Liu X, Peltz G. GSEApy: a comprehensive package for performing gene set enrichment analysis in Python. Bioinformatics (Oxford, England). 2023;39.

111. Doyle JJ, Doyle JL. A rapid DNA isolation procedure for small quantities of fresh leaf tissue. Phytochemical bulletin. 1987.

112. Bolger AM, Lohse M, Usadel B. Trimmomatic: a flexible trimmer for Illumina sequence data. Bioinformatics. 2014;30:2114–2120.

113. Vasimuddin M, Misra S, Li H, Aluru S. Efficient Architecture-Aware Acceleration of BWA-MEM for Multicore Systems. In 2019 IEEE International Parallel and Distributed Processing Symposium (IPDPS); 20-24 May 2019. 2019: 314–324.

114. Li H, Handsaker B, Wysoker A, Fennell T, Ruan J, Homer N, Marth G, Abecasis G, Durbin R. The Sequence Alignment/Map format and SAMtools. Bioinformatics. 2009;25:2078–2079.

115. Danecek P, Auton A, Abecasis G, Albers CA, Banks E, DePristo MA, Handsaker RE, Lunter G, Marth GT, Sherry ST, et al. The variant call format and VCFtools. Bioinformatics. 2011;27:2156–2158.

116. Purcell S, Neale B, Todd-Brown K, Thomas L, Ferreira MA, Bender D, Maller J, Sklar P, de Bakker PI, Daly MJ, Sham PC. PLINK: a tool set for whole-genome association and population-based linkage analyses. Am J Hum Genet. 2007;81:559–575.

117. Lefort V, Desper R, Gascuel O. FastME 2.0: A Comprehensive, Accurate, and Fast Distance-Based Phylogeny Inference Program. Mol Biol Evol. 2015;32:2798–2800.

118. Pritchard JK, Stephens M, Donnelly P. Inference of population structure using multilocus genotype data. Genetics. 2000;155:945–959.

119. Evanno G, Regnaut S, Goudet J. Detecting the number of clusters of individuals using the software STRUCTURE: a simulation study. Mol Ecol. 2005;14:2611–2620.

120. Kopelman NM, Mayzel J, Jakobsson M, Rosenberg NA, Mayrose I. Clumpak: a program for identifying clustering modes and packaging population structure inferences across K. Mol Ecol Resour. 2015;15:1179–1191.

121. Dou T, Wang C, Ma Y, Chen Z, Zhang J, Guo G. CoreSNP: an efficient pipeline for core marker profile selection from genome-wide SNP datasets in crops. BMC Plant Biol. 2023;23:580.

122. DeGiorgio M, Huber CD, Hubisz MJ, Hellmann I, Nielsen R. SweepFinder2: increased sensitivity, robustness and flexibility. Bioinformatics. 2016;32:1895–1897.

123. Quinlan AR, Hall IM. BEDTools: a flexible suite of utilities for comparing genomic features. Bioinformatics. 2010;26:841–842.

124. Camacho C, Coulouris G, Avagyan V, Ma N, Papadopoulos J, Bealer K, Madden TL. BLAST+: architecture and applications. BMC Bioinformatics. 2009;10:421.

125. Zhang Z, Carriero N, Zheng D, Karro J, Harrison PM, Gerstein M. PseudoPipe: an automated pseudogene identification pipeline. Bioinformatics. 2006;22:1437–1439.

