## Supplementary material for "A wild genome of the underutilized legume lablab (*Lablab purpureus*) reveals the genetic basis of domestication": Supp Figure 1

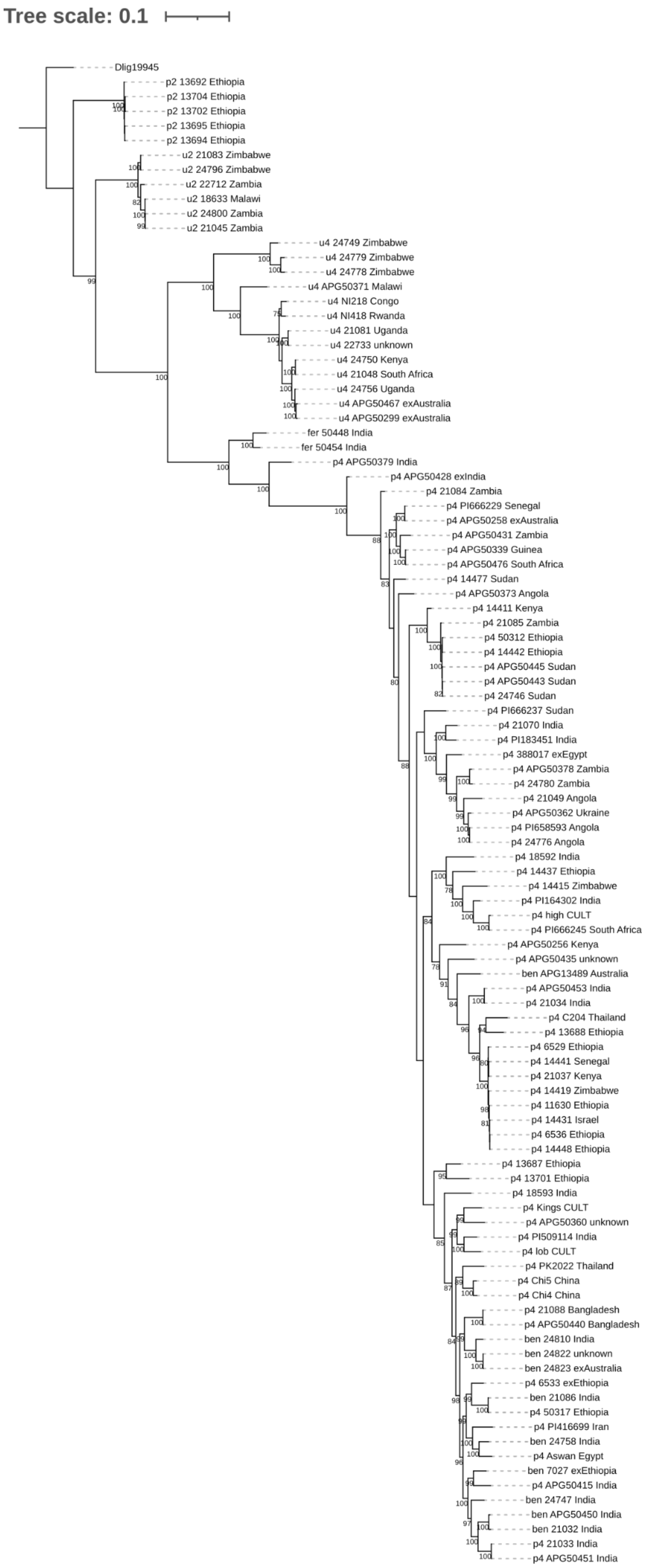


**Fig. S1. Phylogenetic analysis of the full 103 samples.** Samples are labelled as u2, u4, p2, p4 corresponding to wild (subsp. *uncinatus*) two-seeded, wild four-seeded, domesticated (subsp. *purpureus*) two-seeded and domesticated four-seeded, respectively. The NJ tree was computed with 161,064 SNPs (see Methods for details). Branch support on the NJ tree is based on 100 bootstraps, and the tree is rooted on *Dipogon*. Six alleged pairs of samples were included to examine if samples in different seedbanks were indeed identical (see Table S2).
